# *TraCurate*: efficiently curating cell tracks

**DOI:** 10.1101/2020.02.14.936740

**Authors:** S. Wagner, K. Thierbach, T. Zerjatke, I. Glauche, I. Roeder, N. Scherf

**Affiliations:** Institute for Medical Informatics and Biometry, Carl Gustav Carus Faculty of Medicine, TU Dresden, Dresden, Germany; National Center of Tumor Diseases (NCT), Partner Site Dresden, Dresden, Germany; Max Planck Institute for Human Cognitive and Brain Sciences, Leipzig, Germany

## Abstract

TraCurate is an open-source software tool to curate and manually annotate cell tracking data from time-lapse microscopy. Although many studies of cellular behavior require high-quality, long-term observations across generations of cells, automated cell tracking is often imperfect and typically yields fragmented results that still contain many errors. TraCurate provides the functionality for the curation and correction of cell tracking data with minimal user interaction and expenditure of time and supports the extraction of complete cell tracks and cellular genealogies from experimental data. Source code and binary packages for Linux, macOS and Windows are available at https://tracurate.gitlab.io/, as well as all other complementary tools described herein.

## 1 Introduction

Time-lapse microscopy enables the continuous monitoring of individual cells or colonies over time (1). Cell tracking refers to the process in which cellular positions or masks are extracted and linked between observations to obtain a sequential representation of the same object. The required accuracy of cell tracking depends on the biological question at hand. Migration patterns or morphodynamic phenotypes can often be analyzed using automatic tracking algorithms that have only limited accuracy. In contrast, studies of cellular development and behavior often require reliable long-term tracking and unbiased reconstruction of positional and divisional/genealogical information.

Automated tracking algorithms are currently not accurate enough to meet such high standards, as the temporal alignment of cell objects is often error-prone and ambiguous (2). On the other end of the spectrum, completely manual tracking approaches are tedious and labor intensive and thus, severely limit the scale of feasible experiments. TraCurate aims to combine the advantages of automated and manual cell tracking to retrieve accurate, curated cell tracks as well as complete genealogies, and facilitates studies of inter-observer variability. The software and its additional tools provide a complete pipeline to create or import automatically generated tracking data from 2D time-lapse imaging, to manually correct and finally export the curated data sets. TraCurate was designed for but is not limited to, tracking of individual cells in ex vivo assays imaged with widely available standard light microscopes. TraCurate visually presents the images overlaid with the tracking data (either created from our supplementary, minimal-assumption Fiji plugin or imported results from external tracking methods) to allow for computer-assisted post-processing. The user interface is equipped with different, specialized options for user-interaction (called strategies) to allow the user to manually join disconnected tracks of the same object (called tracklets) into cellular genealogies that extend over longer periods and multiple divisions. TraCurate further provides a simple interface to segment, delete, split or merge detected cell masks during curation. As a supplementary tool to TraCurate we provide an R package with a convenient interface for the subsequent statistical analysis of the curated tracking data. This package bridges the gap between our data curation tool and more specialized tools for data analysis.

## 2 Software

TraCurate is implemented in C++11 with Qt5 as a high-level library for the creation of a flexible GUI via QML and abstractions such as different data containers. We rely on the widely used HDF5 library as the basis for the underlying data format as it allows for a compact representation of the tracking data along with additional information for a broad spectrum of potential applications. Data from HDF5 files are represented in-memory using a class hierarchy that mimics the file format itself. The actual image data is an exception here as it is only loaded when the image is displayed in the GUI to keep a low memory profile and enable the responsive editing of high-resolution videos. For data export, we provide the option to save either the complete HDF5 file or only selected parts of the data with a smaller memory footprint (e.g. without the image data). Additionally, we provide an R package to load the curated tracks and image data into common R data structures for subsequent statistical analysis.

### 2.1 Data transformation and format

Over the past two decades a wide range of tools for cell segmentation has been developed (2). Most tools are customized for specific cell types (such as dense cellular monolayers or a few isolated cells), microscopy modalities (e.g. requiring fluorescent labeling of nuclei) or experimental conditions (e.g. high frame rates with considerable spatial overlap between observations at different time points). Our pipeline is designed to be as generic as possible and builds upon minimal assumptions regarding the underlying experiment: We provide an easy-to-use Fiji plugin for 2D cell tracking that creates cell tracklets from segmented images using a simple nearest neighbor tracking as it represents the fewest assumptions about the experiment. The output of this tracking algorithm can be directly imported into TraCurate. Furthermore, by using the open HDF5 format, TraCurate aims to be as compatible as possible with existing, more advanced and specialized tracking algorithms that have also adopted this format, e.g. (3,4). Furthermore, version control of the data format assures compatibility even if it has to be adjusted to novel use cases in the future. A documented specification of the file format simplifies the implementation of interfaces for external developers of other tracking software. At the same time, the HDF5 file type provides the flexibility to store additional meta-data and annotations and offers interfaces in many programming languages.

### 2.2 GUI: graphical user interface

The TraCurate GUI consists of different views: the tracking view, segmentation view and the project view. The main tracking view (see Fig.1) assists the user in connecting tracklets and provides contextual information about the currently selected and hovered cell and the respective tracklet. Typical events that need to be annotated in cell tracking experiments, such as cell division and merging of cells, can easily be assigned to or corrected for a given tracklet, and its status can be specified manually. Furthermore, specialized strategies for fast curation of automatically generated tracks (5) are available to the user. Additional annotations (i.e. nominal classification) can be assigned to objects and tracks to provide further information on the tracking data and enrich the analysis by using annotations based on formal biomedical ontologies (6,7). In the segmentation view, available object outlines can be split and merged if the existing segmentation masks are incorrect. Additionally, TraCurate provides an interactive flood-fill method to add new cell masks that were not automatically segmented. The project view provides general information on the project, such as an overview of the created tracks or the time spent, and allows the specification of user-defined annotations.

**Fig. 1.**
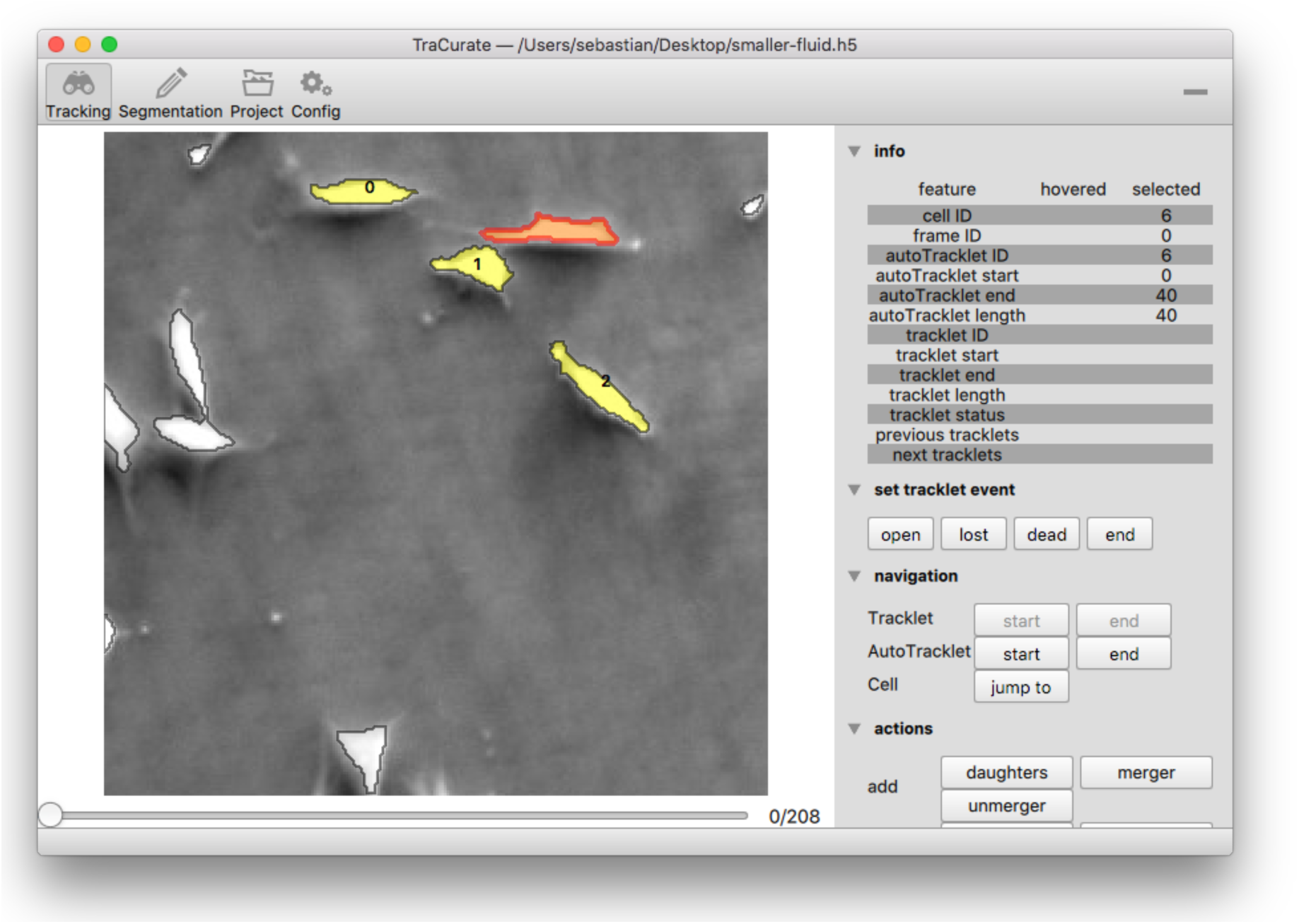
Tracking view of TraCurate. The left part shows the tracking area, in the right menu information on selected and hovered objects is displayed as well as different tools for navigation and editing.

### 2.3 Auxiliary tools and resources

TraCurate aims to integrate into existing workflows as seamlessly as possible and let the user decide which tools to use for segmentation, tracking and downstream analysis. We provide a stand-alone Julia script tc-convert (https://gitlab.com/tracurate/tc-convert) that converts data from various existing formats into the TraCurate HDF5 format. As an example, tc-convert directly supports ilastik HDF5 files (4). With ilastik, the user can train a machine learning classifier with minimal annotations to segment and track cells from time-lapse data. Then manual editing of the automatically generated tracks and curation of cell masks can be done with TraCurate. We further support direct import from the biotracks format, a community-driven standard cell tracking format (8) developed by the Cell Migration Standardisation Organisation (CMSO). Biotracks aims to unify the different cell tracking data formats and itself provides a tool to convert from many different data formats (such as (9)) into the biotracks format. This way, we allow a wide (and growing) variety of tracking tools to be used with TraCurate. If custom tracking tools are used, tc-convert also provides the option to import basic features from a generic CSV format by directly specifying the columns that hold the data.

To also support basic workflows where only cell segmentations (or probability maps) are available, we also supply IJTracker (https://gitlab.com/tracurate/IJTracker_Plugin), a Fiji plugin that implements a simple, unbiased nearest-neighbor approach for automated generation of autotracklets. This plugin records the data in an XML-based format that can be converted to the TraCurate HDF5 format.

To facilitate downstream analysis and visualization of the curated tracking results, we provide tcimport (https://gitlab.com/tracurate/tcimport): a package that allows the user to conveniently import TraCurate HDF5 output into the R language (https://r-project.org), one of the most widely used tools in biomedical data analysis.

The usage of TraCurate and its accompanying tools is described in detail in the TraCurate User Manual, available from the website (https://tracurate.gitlab.io/).

## 3 Comparison to other software

Currently, there exist complementary applications for the (semi-automatic-) tracking of cells. However, to the best of our knowledge, TraCurate is the only multi-platform tool streamlined to creating highly accurate cell shape and tracking data using well-designed interaction strategies to make track curation both easier and faster. Existing tools are either limited to the tracking of cell positions (10), which is a severe limitation for downstream analysis, or to specific cell types and microscopy modalities as they often mix automated tracking with data curation (11). We explicitly separate these two aspects. Consequently, our tool does neither make restrictive assumptions about what is being tracked (e.g. it could be entire organisms) nor about what is being analyzed (e.g. cell shape dynamics). Thus, TraCurate naturally integrates in a variety of experiment and analysis workflows, not forcing the user to apply specific tools for automated cell tracking, downstream visualization, and analysis.

## 4 Case studies

TraCurate (and its preceding development versions) has been used for several applications and has been instrumental for many of our own studies. For example, in (12) we studied the regulation of cell cycle progression on a single cell level. Therefore, we had to track cells over the complete course of the cell cycle and to quantify the intracellular kinetics of several cell cycle regulators continuously. In (13) we studied the effects of heterogeneity in the self-renewal of multipotent hematopoietic progenitor cells. For this purpose, we curated automated but imperfect tracking data to obtain complete cellular genealogies over multiple generations from bright field images. TraCurate was indispensable to obtain highly accurate cell tracks in a semi-automated, time-efficient manner.

